# Structure–function analyses of a keratin heterotypic complex identify specific keratin regions involved in intermediate filament assembly

**DOI:** 10.1101/821090

**Authors:** Chang-Hun Lee, Min-Sung Kim, Shuang Li, Daniel J. Leahy, Pierre A. Coulombe

## Abstract

Intermediate filaments (IFs) provide vital mechanical support in a broad array of cell types. Interference with this role causes cell fragility and accounts for a large number of human diseases. Gaining an understanding IF structure is paramount to understanding their function and designing therapeutic agents for relevant diseases. Here, we report the 2.6 Å resolution crystal structure of a complex of interacting 2B domains of keratin 5 (K5) and K14. K5 and K14 form a long-range, left-handle coiled coil, with participating α-helices aligned in parallel and in register. Follow-up mutagenesis revealed that specific contacts between interacting 2B domains play a crucial role during 10-nm IF assembly, likely at the step of octamer–octamer association. The resulting structural model represents the first atomic-resolution visualization of 2B–2B interactions consistent with the A_22_ dimer alignment and provide insight into the defects introduced by mutations in IF genes associated with human skin diseases.

## INTRODUCTION

Intermediate filaments are major contributors to the homeostasis of cells, organs and tissues, with defined roles in promoting mechanical support, cytoarchitecture, basic cellular processes, and response to stress (Pan et al., 2013; Toivola et al., 2010). Mutations altering the coding sequences of IF genes and proteins, which are conserved and number ~70 in mammalian genomes, are causative for or act as risk factors in the presentation of >90 diseases (Omary et al., 2004; Szeverenyi et al., 2008) (see www.interfil.org database). Yet atomic-level insight into the structure and properties of 10 nm intermediate filaments is lacking at present, though recent progress has been achieved using X-ray crystallography (Guzenko et al., 2017) and cryo-electron tomography (Turgay et al., 2017). Understanding the structure of IFs at high resolution is essential to fully appreciate their cellular roles, how these are disrupted in disease, and for drug design and development.

Early efforts to study the core structure of 10 nm IFs entailed the use of nearest-neighbor analyses using crosslinking agents coupled with site-directed mutagenesis. Studies focused on vimentin (type III IF) and keratins (type I and II IFs) revealed four distinct alignments of dimers in the polymer lattice, A_11_, A_22_, A_12_, and A_CN_ (Figure S1) (Geisler et al., 1992; Steinert et al., 1993a; Steinert et al., 1993b). Using homology modeling and mutagenesis, Bernot et al. (Bernot et al., 2005) provided evidence that skin keratins mainly use the 1B domain to form a stable anti-parallel tetramer with A_11_ alignment (see (Geisler et al., 1992)). Moreover, there is crystallographic insight for the contacts responsible for the A_11_ (Aziz et al., 2012, Eldirany et al., 2019) or A_CN_ alignments (Strelkov et al., 2004). While there is structural information for coiled coils comprising the 2B domain for several IFs (Bunick and Milstone, 2017; Chernyatina et al., 2012; Lee et al., 2012; Nicolet et al., 2010; Strelkov et al., 2001; Strelkov et al., 2002; Strelkov et al., 2004), high resolution insight into the structural determinants for the A_22_ and A_12_ configurations is still lacking.

We previously determined the 3.0 Å crystal structure of the interacting 2B domains of keratin 5 (K5) and keratin 14 (K14) (Lee et al., 2012). K5 and K14 respectively are type I and type II IF sequences that interact in an obligatory fashion to form 10 nm filaments in progenitor basal keratinocytes of epidermis and related epithelia (Fuchs, 1995; Nelson and Sun, 1983). K5-K14 IFs serve as anchors for tissue architecture in epidermis and are mutated and defective in epidermolysis bullosa simplex, a rare condition in which basal keratinocytes are fragile and prone to rupture upon exposure to frictional stresses (Coulombe et al., 1991; Coulombe and Lee, 2012). The 2B region (~117 residues) features long range, coiled-coil forming heptad repeats and represents a prominent segment in the highly-conserved and functionally essential central α-helical rod domain shared by all IF proteins (Guzenko et al., 2017; Lee et al., 2012). While the K5-K14 co-crystal provided insight into heteromeric IF coiled-coils, the unexpected presence of a homotypic, transdimer disulfide bond mediated by residue cysteine 367 in human K14 had a major impact on contacts made in the crystal lattice and limited its significance for the interactions between heterodimer and, higher-order subunits within filaments (Lee et al., 2012). The same outcome occured when crystallizing the corresponding segments from K1-K10 (Bunick and Milstone, 2017), characteristic of differentiating keratinocytes in epidermis and related epithelia (Fuchs, 1995). Here, we report on insight gained from the crystal structure of the interacting 2B domains of K5-K14 in which disulfide bond formation is lost due to a Cys-Ala mutation at position 367 in K14.

## RESULTS

### Structure of K5wt/K14 C367A 2B Domains

We reported on the structure of 2B domain from keratin K5/K14, which forms a heterodimeric coiled coil structure with a homotypic trans-dimer disulfide bond linking adjacent dimers (Lee et al., 2012). In this structure, all contact areas between K5/K14 dimers in the crystal are buried in less than 500 Å^2^, suggesting that they may not occur in native keratin IFs. Available evidence suggests that the disulfide bond(s) mediated by K14’s Cys367 likely occurs between rather than within 10 nm filaments (Lee et al., 2012).

To identify protein-protein interactions between the 2B domains of K5 and K14 that may be important for keratin assembly and/or IF structure, we mutated K14’s cysteine 367 residue to alanine (C367A) and set up crystallization trials with K5. This resulted in a new crystal form that diffracted X-rays to 2.6 Å resolution (Table 1). The K5wt-K14C367A structure was determined by molecular replacement and encompasses 95 amino acids in K14 (residues 327-421) and 98 amino acids in K5 (residues 379-476). The overall structure and relative orientation of coiled coil helices in the K5wt-K14C367A complex are highly similar to that reported for K5wt-K14wt 2B heterodimers (Protein Data Bank (PDB) ID: 3TNU) with an r.m.s deviation of Cα positions of 0.959 Å over 83 amino acids in K14 (residues 336-418) and 86 amino acids in K5 (residues 389-475) (Figure 1A).

**Table 1.** Statistics of X-ray data collection and refinement

| **Data collection** |  |
| --- | --- |
| Radiation Source | APS GM-CA |
| Wavelength (Å) | 0.97838 |
| **Data processing** |  |
| Space group | P41 |
| Copies / ASU | 1 |
| Cell dimensions |  |
| a, b, c (Å) | 56,15, 56,15, 222,05 |
| $\alpha$, $\beta$, $\gamma$ (°) | 90, 90, 90 |
| Resolution range (Å)<sup>a</sup> | 37.38-2.60 (2.69-2.60) |
| Number of reflections (unique) | 20213 (1888) |
| Multiplicity | 4.6 (2.6) |
| Completeness (%) | 96.08(89.86) |
| R<sub>mean</sub> | 8.5 (34.8) |
| I/ $\sigma$ (I) | 8.91 (2.66) |
| CC ½ | 99.7 (85.7) |
| **Refinement** |  |
| Resolution (Å) | 37.38-2.60 (2.69-2.60) |
| R<sub>work</sub> /R<sub>free</sub> (%) | 20.1 / 24.6 |
| No. protein atoms | 3079 |
| Mean B –value proteins atoms | 135.50 |
| R.m.s deviations |  |
| Bond lengths (Å) | 0.006 |
| Bond angles (°) | 0.76 |
| Ramachandran statistics (%)<sup>b</sup> |  |
| Most favored regions | 96 |
| Additional allowed regions | 4 |
| Disallowed regions | 0 |

<sup>a</sup> Highest resolution shell is shown in parentheses.
<sup>b</sup> As defined by the validation suite MolProbity.

**Figure 1.**
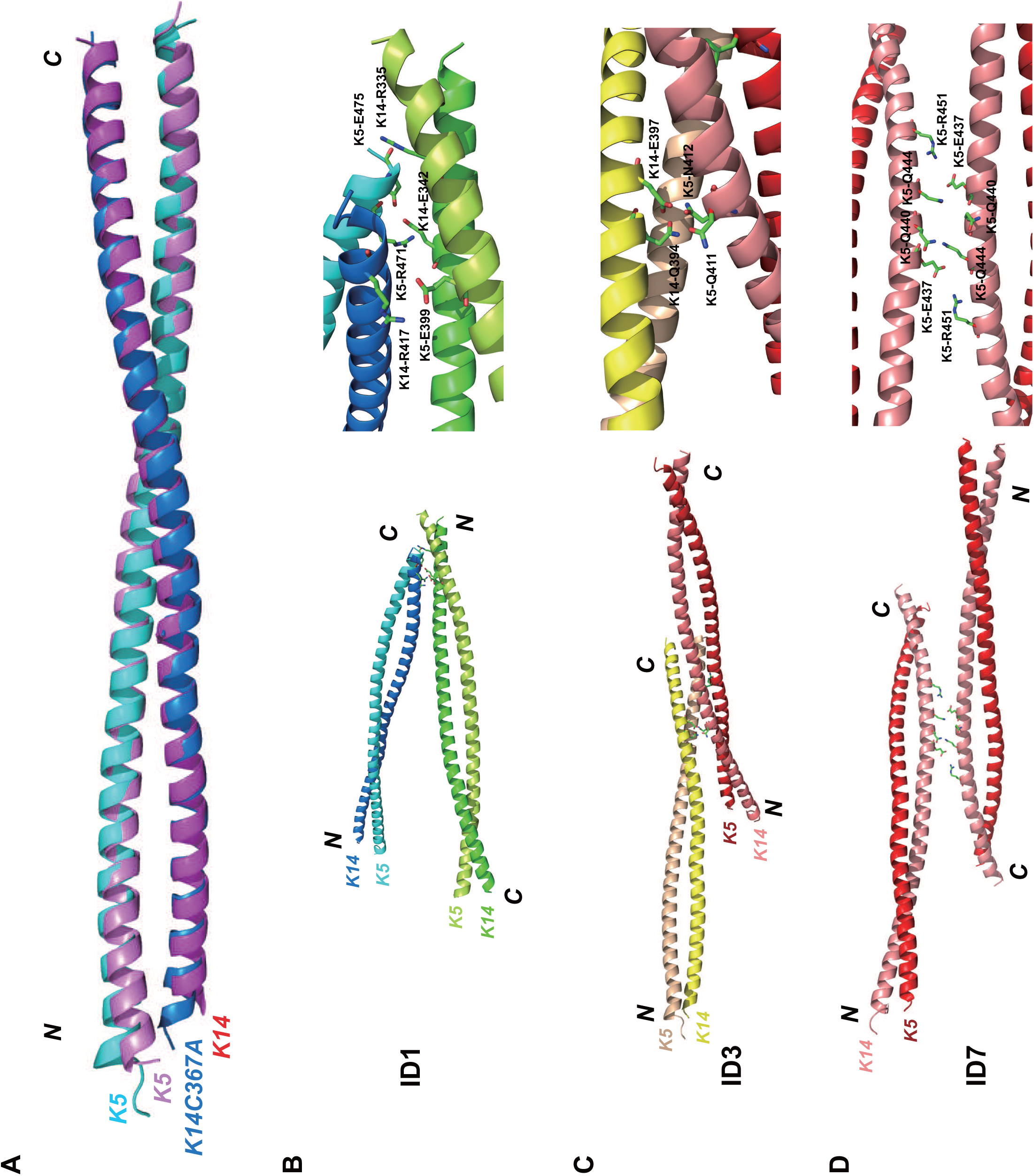
Modeling 2B-2B interactions in IFs from the crystal structure of K5wt/K14-C367A. (A) Two K5/K14 heterodimer models are shown superimposed. (B-D) Three interaction interfaces occurring in the crystal structure. Left panels show interacting coiled-coil dimers. Right panels show close-up view of key interacting residues between dimers. (B) ID1 shows anti-parallel interactions between dimers. (C) ID3 shows parallel interactions between dimers. (C) ID7, corresponding to the asymmetric unit in crystal, shows anti-parallel interactions between dimers.

The K5wt-K14C367A mutant crystal exhibits multiple contacts between dimers not present in the K5wt-K14wt structure (Figure S2 and Table S1). PISA (Protein Interfaces, Surfaces, and Assemblies) analysis identified two main regions of inter-dimer (ID) contacts (Table S2). One contact, designated ID1 (Figures 1B and S3), involves antiparallel interactions between the N-terminal part of a K5/K14 heterodimer (K14 residues R335, M338, Q339, E342, L345, Q346, L349, S350, K352, A353, N357; K5 residues M389, M392, R395, L396, E399) and the C-terminal part of a neighboring heterodimer (K14 residues K399, D403, R407, Q410, E411, T414, Y415, R417, L418; K5 residues R471, L474, E475). Another ID contact, ID3 (Figure 1C), involves parallel interactions between the N-terminus of a K5/K14 dimer (K14 residues Q394, E397; K5 residues Q411, N412) and the proximal middle part of another K14/K5 dimer (K14 residues R407, R365, Y366, Q372; K5 residues E422, D464, R471). ID1 and ID3 bury 805.7 Å^2^ and 760.9 Å^2^, respectively, and do not overlap. The residues involved in these contacts are highly conserved in the keratin superfamily (Figure S4) and may have important and conserved roles during keratin IF assembly. Additional ID contacts occur in the K5wt-K14C367A mutant crystal lattice but the surface areas involved are < 200 Å^2^ and unlikely to play a major role during keratin assembly. PISA also showed that pairs of dimers make an ID contact, ID7, with a buried surface area of 215.3 Å^2^ in the asymmetric crystal unit (Figure 1D and Table S1). The ID7 contact is intriguing since it involves the middle segment of the K5 coil and involves symmetric interactions between opposing E437, Q440, Q444 and R451 residues, and was used as a reference in subsequent analyses (Table S2).

### ID1 Contact-deficient Keratins Cannot Form IF Networks in Transfected Cells

We hypothesized that the ID1 and ID3 contacts may reflect molecular interactions important for 10 nm filament assembly and structure. To test this possibility, we mutated key residues in each of the ID1, ID3 and ID7 interfaces (see Figure 1B-1D, Figure S4 and Table S2) to alanine residues and transfected the resulting full-length K5 and K14 constructs into NIH 3T3 fibroblasts. In selecting residues for mutagenesis we purposely avoided the C-terminus of K5 and K14 2B domains since they are already known to be important for IF assembly and/or are mutated in EBS (Gu and Coulombe, 2005; Gu et al., 2002; Letai et al., 1993; Yasukawa et al., 2002). Accordingly, we chose K14-R335, K14-E342, K14-Q346 and K5-E399 to generate ID1-deficient mutants, and K5-Q411, K5-N412, K5-E422, K14-R365, K14-Y366 and K14-Q372 to generate ID3-deficient mutants. An ID7-deficient mutant was made by changing K5-E437, K5-Q440 and K5-Q444 to alanine (see Methods and Table S2). NIH 3T3 cells do not express keratin proteins, thus providing an ideal live cell setting in which to test for de novo keratin assembly (Paladini et al., 1996). In transfected cells expressing wildtype K5 and K14, keratin IF networks are well-developed with fibers radiating across the cytoplasm, as is typical of skin keratinocytes (Figures 2A and 2B). Only a small fraction (<10%) of these cells showed keratin aggregation. A similar outcome occurred in cells expressing the mutant pairings compromising the ID3 or ID7 contact-interfaces (Figures 2A and 2B). By contrast, >90% of NIH 3T3 cells expressing the ID1-contact-deficient keratin pair showed keratin-positive aggregates (Figures 2A and 2B). A small fraction of these cells showed keratin IF networks that were very thin and sparse in the cytoplasm. Quantification confirmed that 93.2±1.1%, 91.7±1.3%, and 92.6±2.5% of transfected cells showed well-developed keratin networks when expressing the wildtype, ID3 mutant and ID7 mutant pairings, respectively, but only 9.2±2.6% for ID1 mutant-expressing cells (Figure 2B). This non-epithelial cell transfection assay thus provides evidence that ID1 contacts between the 2B domains of K5 and K14 are key to forming keratin IF networks in vivo.

**Figure 2.**
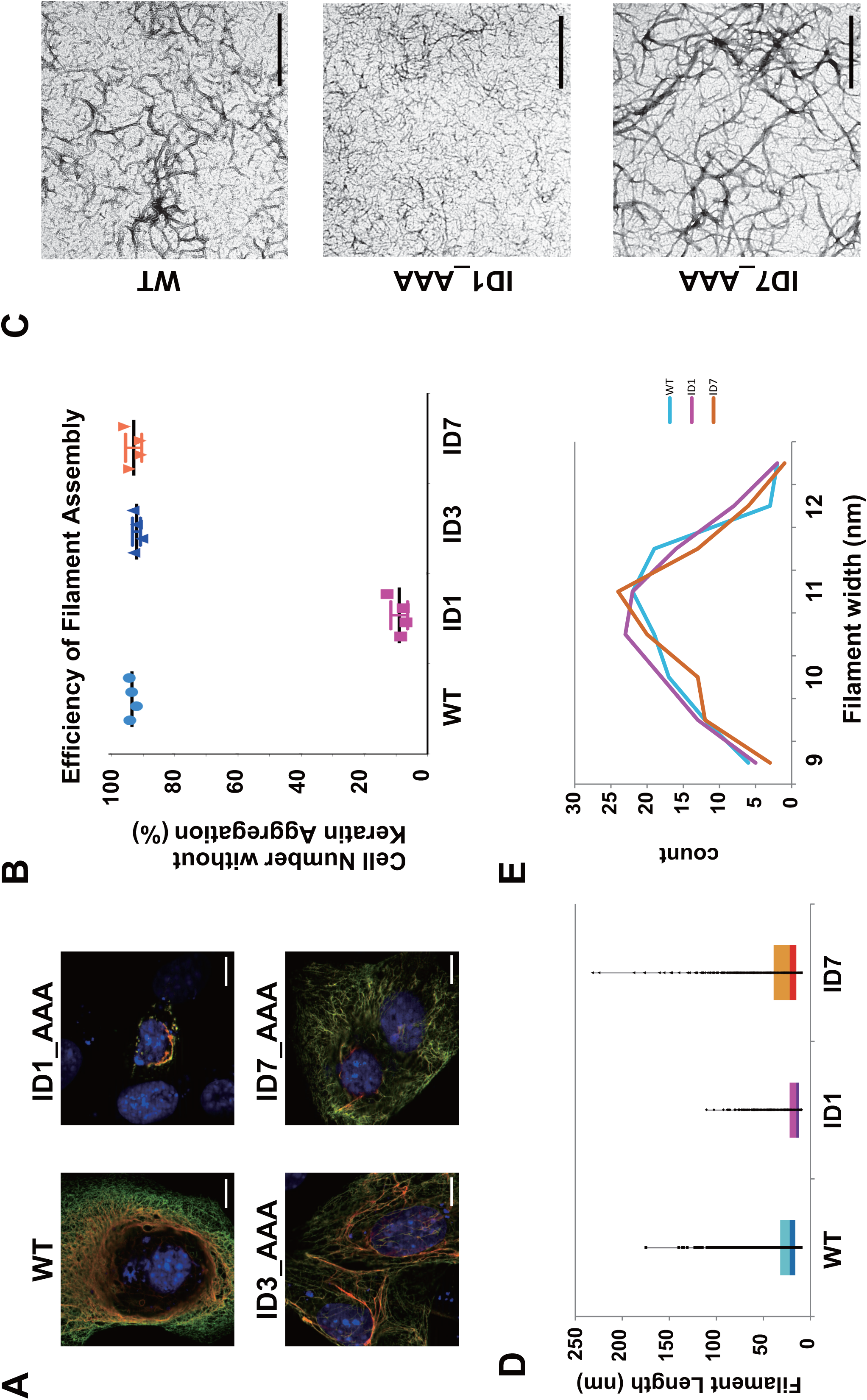
Functional assays with K5 and K14 mutant pairs show that ID1 contacts are key to IF assembly. (A) Representative images of NIH-3T3 cells transfected with wildtype K5/K14 or mutant K5/K14 pairings (K5, green; K14, red; nuclei (DNA), blue). Bar, 10 μm. (B) Quantitation of transfected cells showing normal-looking keratin filament networks (four independent trials for each combination tested). Imaged cells (n = 100 per sample) were randomly selected. (C) Micrographs of wildtype and mutant keratin assemblies (100 μg/ml) subjected to uranyl acetate staining. The images show positively-stained filaments. Bar, 200 nm. (D) Probability of observing long filaments in positively-stained TEM images, calculated as the (pixel) ratio of long vs. short filaments (ImageJ). (E) Distribution of filament width in negatively-stained TEM images. For each sample, 100 filaments were selected and measured using ImageJ; relative ratios are reported. The color code used (B, D, E) is: cyan, wildtype; magenta, ID1 mutant; orange, ID7 mutant. Wildtype and mutant keratins form filaments having similar width.

### ID1-contact-deficient Keratins Form Abnormally Short IFs

We next examined the role of the ID1 contact interface during 10 nm IF assembly using purified recombinant proteins in vitro (Coulombe and Fuchs, 1990; Lee and Coulombe, 2009). Assemblies were viewed by negative-stain transmission electron microscopy (EM). As expected, the wildtype K5-K14 pairing showed a high density of filaments that were long with a regular and smooth outline (Figure 2C, upper row). A similar outcome was observed with the ID7 mutant pairing, but not with the ID1 pairing (Figure 2C). While challenging owing to the high density of filaments, quantitation of the frequency of long filaments confirmed that the ID1 mutant pairing showed a markedly reduced fraction of long filaments (Figure 2D). Higher magnification views showed that the width of individual IFs was the largely the same among wildtype, ID1 mutant and ID7 mutant pairings (Figure 2 C) and was ~10 nm (Figure 2E). These EM observations from purified proteins are consistent with and extend the NIH 3T3 cell transfection data and suggest that the ID1 and ID7 interfaces do not play a major role in the initial phase of 10 nm filament assembly (lateral growth) but that ID1 contacts are crucially important during the elongation phase of this process.

## DISCUSSION

### A Role for ID1 Contacts during IF Assembly

We report here on a crystal structure of the interacting 2B domains of K5 and K14 in which the potential for the formation of the trans-dimer, homotypic disulfide bond was eliminated. The resulting structure confirms the K5-K14 coiled-coil dimer structure originally reported (Lee et al., 2012) but reveals a dramatically different set of interactions between dimers in the crystal lattice. Follow-up analyses involving mutagenesis revealed a role for a specific region of the K5 and K14 2B domains in the inter-subunit interactions and 10 nm IF assembly. This specific interface, ID1, buries 805.7 Å^2^ of surface area and involves residues 385-403 in K5 (chain 1) with 331-361 in K14 (chain 1’) and 467-476 (chain 2) in K5 with 399-421 (chain 2’) in K14, which are exposed on the heterodimer surface and interact antiparallel with a 39.4° angle between the suprahelical axes of dimers. The outcome of 10 nm filament reconstitution assays in vitro implies that these contacts are forming as short filaments anneal into longer polymer strands. The alanine mutations we introduced to disrupt the ID1 interface may themselves have a deleterious impact of keratin IF assembly, an unlikely prospect that can best addressed by additional structural studies. Short of this possibility, the ID1 contact interface is poised to play a role in guiding IF octamer assembly through 2B-2B interactions, as is further discussed below.

The C-termini of both K5 and K14 rods features amino acids that are highly conserved across the IF protein superfamily, notably the signature “TYRR(K)LLEGE” motif (Parry, 2005). K14 residues T414, Y415, R417 and L418, and K5 residues R471, L474 and E475 (Figures 1B and S3) participate in the nucleation and stabilization of coiled coil dimers (i.e., act as a “trigger motif”) via a network of ionic and hydrophobic interactions (Wu et al., 2000). Missense alleles affecting residues mapping to this conserved motif in each of K5 and K14 act dominantly to cause generally severe variants of EBS (Coulombe and Lee, 2012; Omary et al., 2004; Szeverenyi et al., 2008). The side chains of K14-E411, K14-R417 and K5-R471 are exposed at the surface of the coiled coil, suggesting a role during assembly beyond dimer formation. By using mutagenesis coupled to functional assays, Wu et al. (Wu et al., 2000) showed that Glu106 (which corresponds to K14-E411 and K5-E466) has a defining role in stabilizing both coiled-coil structure and the A_22_ interaction (Figure S1). The structural model below explains how mutation of this residue can affect the stability of tetramers with the A_22_ configuration, which is likely to play a key role during the annealing of oligomers or short filaments.

The relevance of the newly identified ID1 interface to 10 nm IF assembly and function is strongly supported by the occurrence of several variants causative for EBS, including K14-E411 (mutated to K in EBS-Dowling-Meara (DM), which is severe, and in EBS-localized), K14-R417 (mutated to P in EBS-DM), K14-L418 (to V in EBS-generalized), K5-R471 (to C in EBS-localized), K5-E475 (to K or G in EBS-DM) and K5-G476 (to D in EBS-localized or EBS-EPPK). These missense alleles act dominantly and tend to be associated with more severe forms of EBS. The similarities between the assembly properties of EBS-causing mutants and those of ID1 interface mutants add to the notion that the ID1 contact serves as a major stabilizing step during keratin IF assembly. Mutagenesis studies performed on other keratins or types of intermediate filament proteins now appears as a worthwhile endeavor to determine whether the ID1 contact is of general importance during IF assembly.

The parallel inter-dimer contact corresponding to the ID3 interface also contains amino acid residues related to EBS, including K5-K404 (mutated to E in EBS-localized), K14-E411 (to K in EBS-DM and EBS-generalized), K5-R471 (to C in EBS-generalized) and K5-E475 (to K or G in EBS-DM) (see www.interfil.org). In this instance, again, analysis of keratin crosslinks (Wu et al., 2000) identified pairings comprised of K5-K404 and K5-443, K5-K404 and K5-441, and K5-K405 and K5-K443. These crosslinks are consistent with a model of parallel alignment of interacting 2B domains from K5, and are accounted for in the ID3 interface. Though a role for ID3 is not supported by our mutagenesis data, these findings suggests that parallel alignment of interacting 2B subdomains also occurs within the IF.

### Implications for Keratin Filament Assembly

Our current understanding of many of the molecular interactions occurring during keratin heteropolymer assembly lags behind that of other IF proteins, notably vimentin and lamins (Guzenko et al., 2017; Herrmann et al., 2007). Based on evidence reported here, we propose a model for the interactions occurring between larger-sized subunits during keratin assembly.

As for all other IFs, polymerization of keratin IFs begins with the formation of coiled-coil dimers, heterodimers in the case of keratin (Coulombe and Fuchs, 1990; Hatzfeld and Weber, 1990; Steinert, 1990), with participating type I and II IF monomers aligned in parallel and in register (Hatzfeld and Weber, 1990). Heterodimers interact along their lateral surfaces in a staggered and antiparallel fashion to form structurally symmetric tetramers (Aebi, 1988; Guzenko et al., 2017; Herrmann et al., 2007). For cytoplasmic IFs, specifically, the lateral contacts productive for 10-nm filament formation occur between the 1B subdomains of participating dimers, oriented anti-parallel (Bernot et al., 2005; Eldirany et al., 2019; Geisler et al., 1992), generating a conformation known as the “A_11_ alignment” (Figure S1; (Steinert et al., 1993a)). Tetramers interact to form octamer subunits (Sokolova et al., 2006) via lateral contacts involving 1B-1B and 2B-2B pairings. At this stage, the interacting 2B domains are by necessity oriented in parallel (Figure 3). SAXS (Small Angle X-ray Scattering) data and modeling (Sokolova et al., 2006) suggested an octamer model that involves the A_11_ alignment (Figure S1) and accommodates a “unit-length filament” (ULF) intermediate. Three additional studies suggested structural models of “1B tetramers” involving 1B-1B antiparallel interactions via the A_11_ alignment (Aziz et al., 2012; Pang et al., 2018, Eldirany et al., 2019). Aziz et al.’s model of tetramer structure (Aziz et al., 2012) is supported by crystallographic data and site-directed spin labeling electron paramagnetic resonance.

**Figure 3.**
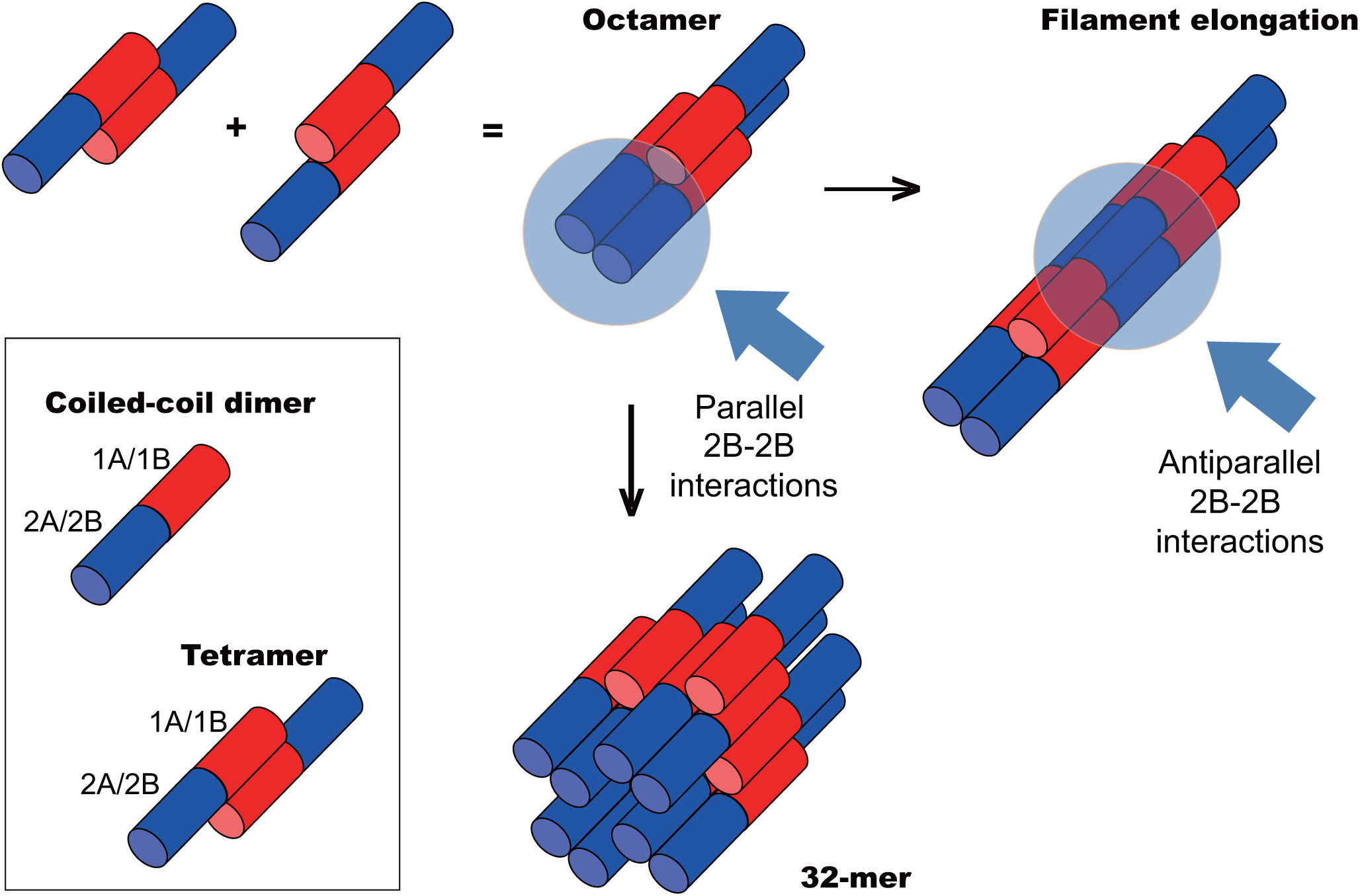
Model for subunit interactions during keratin assembly. Keratin heterodimers are depicted as bi-colored rods to reflect structural asymmetry. The N-terminal half (red) represents coil 1 segment (1A, 1B subdomains). The C-terminal half (blue) represents coil 2 segment (2A, 2B subdomains) of the rod (Fig. S1). The head and tail domains, of undefined structure, are not shown. Tetramer and octamer units are presented based on the 1B-1B interaction model (A_11_) (see text). Anti-parallel 1B-1B interactions and parallel 2B-2B interactions occur as subunits interact laterally to yield a 32-mer assembly with a diameter of 10-nm. Anti-parallel 2B-2B interaction (A_22_) occur when octamer intermediates or 32-mer complexes interact end-to end, thus lengthening (elongating) the polymer strand.

Mass determination based on electron microscopic images suggests that reconstituted keratin IFs are comprised of 32 polypeptide chains or four octamers in cross-section (Aebi, 1988), implying that octamers interact both laterally and end-to-end in the 10-nm filament lattice. We propose that assembly steps beyond octamer formation entail 2B-2B interactions in two complementary ways. End-to-end annealing of octamers necessitates interactions between participating 2B subdomains oriented antiparallel (Figure 3). Our data suggests that those interactions involve the ID1 interface and are consistent with the A_22_ alignment (Figure S1). By contrast, lateral interactions between octamers must involve the docking of 2B subdomains oriented in parallel (Figure 3) – further studies are needed to identify the contacts involved. This model rationalizes why ID1 interface mutants form only short IFs in vitro. In conclusion, we present evidence that direct interactions involving specific regions in the coil 2 segment of both K5 and K14 (the ID1 contact) play a key role during IF formation, most likely at a step after octamer assembly. Our structural model is the first high resolution depiction of 2B-2B interactions that may reflect Steinert’s A_22_ alignment between subunits. Our findings further our understanding of the molecular mechanism underlying IF assembly and the defects introduced by mutations in disorders such as epidermolysis bullosa simplex.

## ACKNOWLEDGMENTS

The authors are grateful to members of the Coulombe and Leahy laboratories for support. The work was supported by grant AR042047 from the National Institutes of Health to P.A.C. and award RR160023 from the Cancer Prevention and Research Institute of Texas to D.J.L..

## AUTHOR CONTRIBUTIONS

C.H.L. and P.A.C. designed the study, interpreted the data, and wrote the manuscript. M.S.K. and D.J.L. provided crystallographic expertise, and M.S.K. helped with data interpretation and presentation. S.L. conducted the cell transfection assays and some of the in vitro assembly assays. All other experiments were carried out by C.H.L and M.S.K.

## STAR METHODS

Detailed methods are provided in the online version of this paper and include the following:

- KEY RESOURCES TABLE
- CONTACT FOR REAGENTS AND RESOURCE SHARING
- EXPERIMENTAL MODEL AND SUBJECT DETAILS
  - NIH-3T3 Cell Culture
- METHOD DETAILS
  - Protein Purification
  - Crystallization
  - Structure determination and analyses
  - Site-directed Mutagenesis to disrupt 2B-2B subdomain interactions
  - Evaluation of keratin filament assembly in cultured cells
  - Evaluation of keratin filament assembly in vitro
- DATA AND SOFTWARE AVAILABILITY

### Key Resources Table

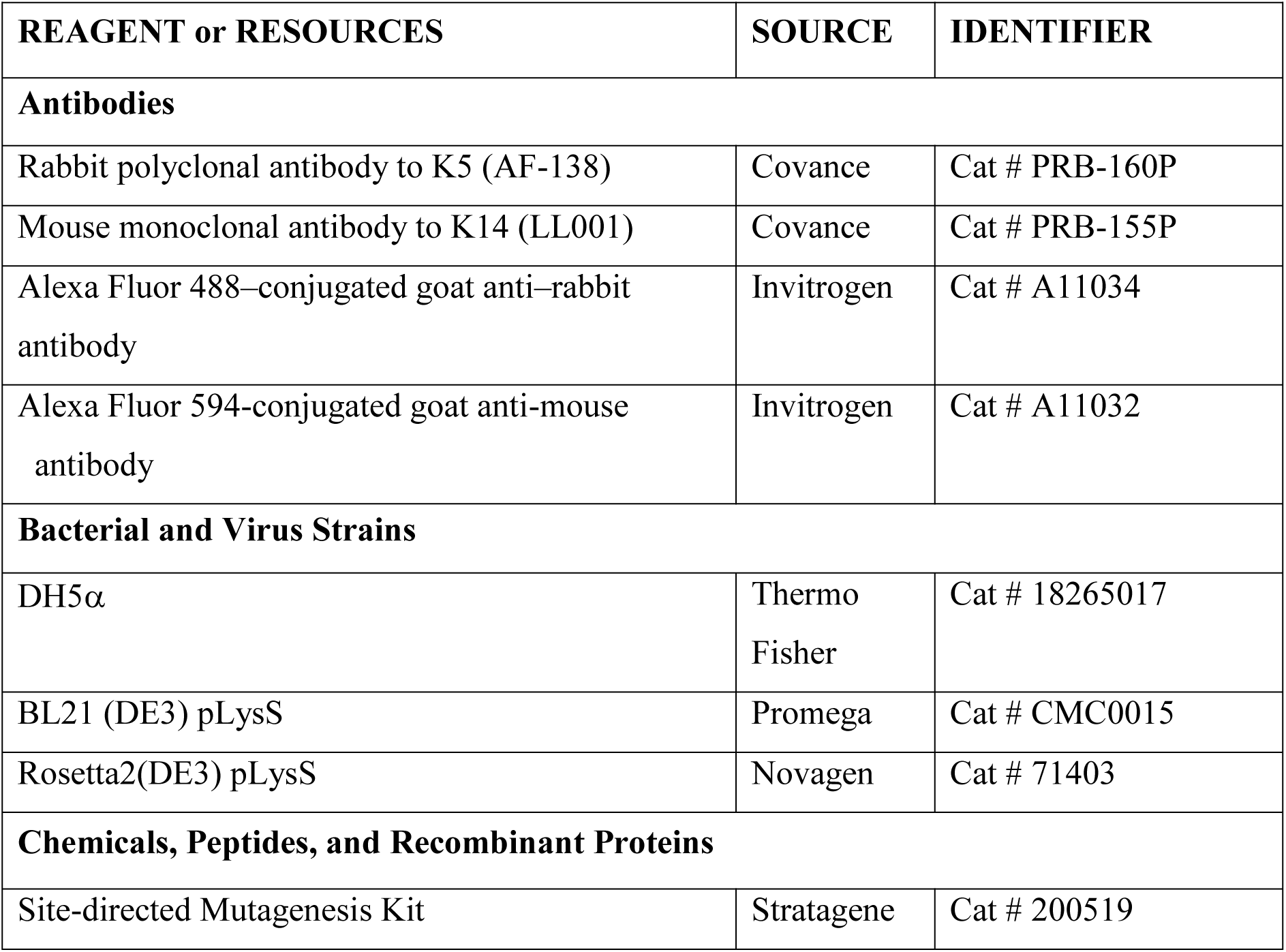
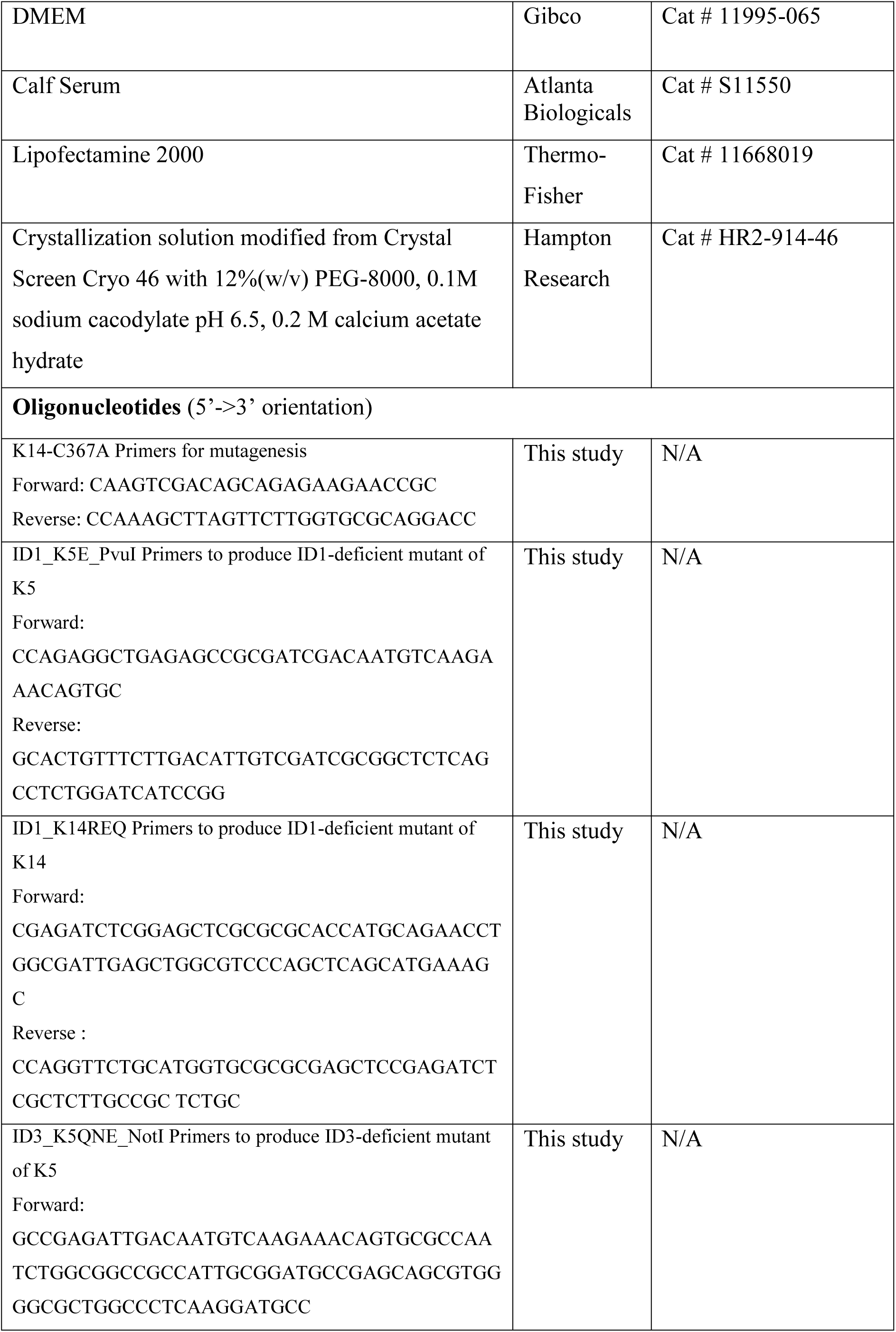
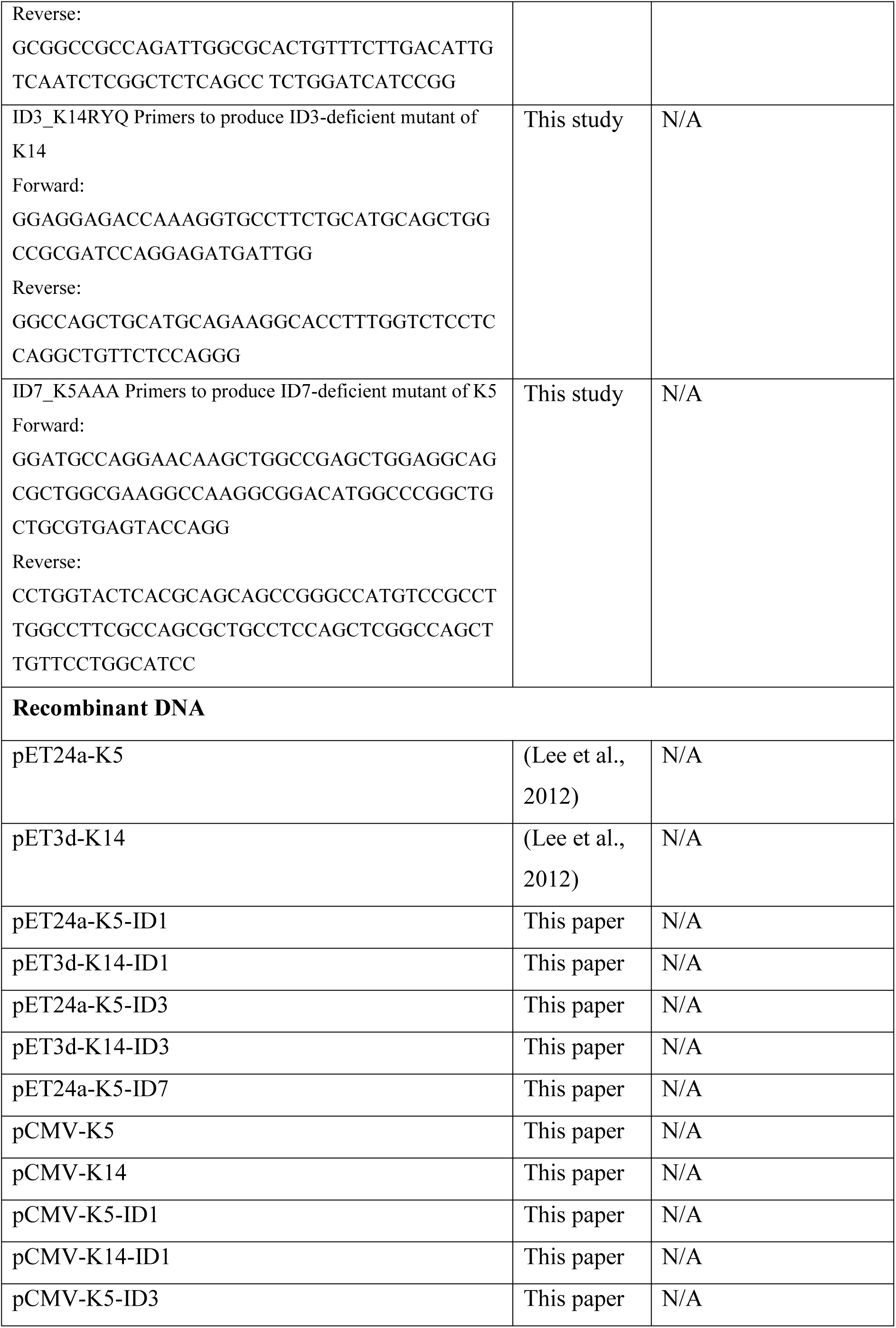
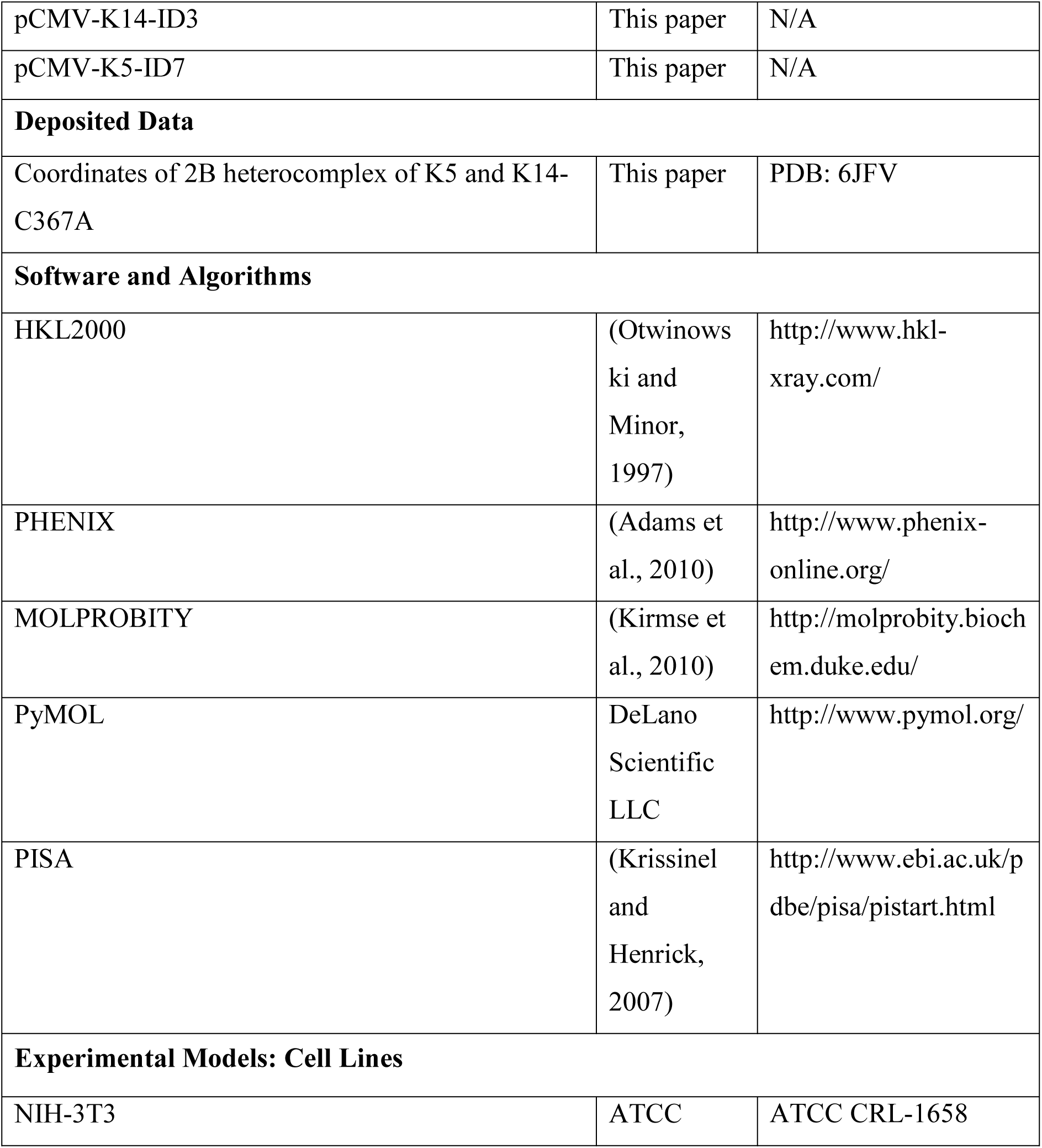

## CONTACT FOR REAGENT AND RESOURCE SHARING

Further information and requests for resources and reagents should be directed to and will be fulfilled by the Lead Contact, Pierre A. Coulombe.

## EXPERIMENTAL MODEL AND SUBJECT DETAILS

NIH-3T3 cell were cultured in DMEM containing 2mM glutamine and 10% calf serum (VWR).

## METHODS DETAILS

### Protein Purification

Plasmids for bacterial expression of the 2B subdomains of human K5 (residues 350-478) and K14 (residues 295-472) and protocols for protein purification are described (Lee et al., 2012). K14’s codon 367 was mutated from cysteine to alanine (C367A) using PCR-directed mutagenesis (Lee and Coulombe, 2009; Lee et al., 2012) with oligonucleotide primers (K14-C367A) listed in the Key Resource Table.

### Crystallization

Crystals of the K5–K14-Cys367A 2B heterocomplex were grown at 20 °C via the hanging-drop vapor diffusion method by mixing 2 μl of protein solution (10 mg ml−1) with 2 μl of reservoir solution (12% (w/v) PEG-8000, 0.1M sodium cacodylate pH 6.5, 0.2 M calcium acetate hydrate) (Hampton Research). Amorphous crystals were crushed and seeded into the same mixture to grow larger crystals (P4 space group) that were frozen (liquid nitrogen) and sent for data collection to the Advanced Photon Source beamline at Argonne National Laboratory.

### Structure determination and analyses

Diffraction data were processed with the software HKL2000. The structure was determined by molecular replacement using Phaser-MR in PHENIX with K5 L2-2B/K14 L2-2B dimer structure (PDB:3TNU) as search model. The initial model was refined using PHENIX and validated using Molprobity (Chen et al., 2010). Interactions between dimers were assessed with PISA (http://www.ebi.ac.uk/pdbe/pisa/pistart.html). Molecular images were generated using the PyMol Molecular Graphics System, Version 2.2 (Schrödinger, LLC).

### Site-directed Mutagenesis to disrupt 2B-2B subdomain interactions

Amino acids potentially interacting in the context of 2B-2B interfaces were identified in the crystal lattice and replaced with alanine (see Tables S2 and S3) via site-directed mutagenesis (Stratagene). The mutants generated and oligonucleotide probes used for PCR mutagenesis are listed in Key Resource Table. All target mutation sites and coding sequences were verified by DNA sequencing (Macrogen USA, MD). Relevant DNA fragments were cloned into pET24a (K5 derivatives) and pET3d (K14 derivatives) plasmids for bacterial expression, or into pCMV vectors (both K5 and K14 derivatives) for mammalian cell expression.

### Evaluation of keratin filament assembly in cultured cells

NIH-3T3 cells were grown on coverslips in DMEM containing 2mM glutamine and 10% calf serum (VWR). DNAs of full length K5 and K14 (wildtype or mutant) were dual-transfected using lipofectamine 2000 (Thermo-Fisher) and cultured for 72h. Cells were processed for indirect immunofluorescence (Lee and Coulombe, 2009) and imaged using an AxioObserver inverted microscope with Apotome 2 optical sectioning module and AxioCam MRm camera (Carl Zeiss). Primary antibodies used were a rabbit polyclonal antibody to K5 (AF-138; Covance; 1:1,000) and a mouse monoclonal antibody to K14 (LL001; Covance; 1:1,000). Secondary antibodies included an Alexa Fluor 488–conjugated goat anti–rabbit and Alexa Fluor 594-conjugated goat anti-mouse (Invitrogen; 1:1,000). Data from keratin-stained images were analyzed using Microsoft Excel.

### Evaluation of keratin filament assembly in vitro

pET24a-K5 and pET3d-K14 constructs (wildtype or mutant) were transformed into the E. coli strain BL21 (DE3) pLysS to overexpress proteins as inclusion bodies (Coulombe and Fuchs, 1990). Recombinant proteins were purified as described (Ma et al., 2001). Full length K5 and K14 proteins were mixed (800 μg/ml) and subjected to MonoQ purification to recover heterotypic complexes in a 1:1 molar ratio. Heterotypic complexes (100-400 μg/ml) were subjected to polymerization by overnight dialysis at 4°C into filament assembly buffer (5 mM Tris-HCl, pH7.4; see (Coulombe and Fuchs, 1990; Lee and Coulombe, 2009; Ma et al., 2001). Filament ultrastructure was examined by negative staining (1% uranyl acetate) and transmission electron microscopy (Philips Bio-Twin CM120. FEI Company). Size information from images was computed using ImageJ.

## DATA AND SOFTWARE AVAILABILITY

Coordinates and structure factors for the K5–K14-Cys367A 2B heteromeric complex were deposited in the Protein Data Bank with code 6JFV.

## SUPPLEMENTAL INFORMATION

Supplemental Information is provided at “Lee et al. Supplemental Information.pdf”. Materials included in this Supplemental Information are as followed:

Figure S1. Schematic depicting the four modes of interaction between dimers of IF proteins in assembled intermediate filaments.

Figure S2. Comparison of crystal contacts made by wildtype K5/K14 2B complexes (left) vs. K5/K14-C367A mutant 2B complexes (right).

Figure S3. Electron density map of the ID1 contact in K5/K14-C367A mutant 2B complexes.

Figure S4. Sequence alignment of type I and type II keratins in the 2B region of the central rod domain.

Table S1. List of ID contacts found in the K5/K14-C367A mutant 2B complexes.

Table S2. Summary of PISA analysis and mutated residues to produce ID deficiencies.

Table S3. Oligonucleotide primers used for mutagenesis to produce ID-deficient mutants.

