## Supplemental Figs S1-S3 and Tables S1-S3 for "Structure–function analyses of a keratin heterotypic complex identify specific keratin regions involved in intermediate filament assembly"

**List of elements included in this <u>Supplemental Information</u> section:**

**Figure S1. Four modes of interaction between dimers of IF proteins.**

**Figure S2. Crystal contacts of K5/K14 wildtype 2B complex and of K5/K14-C367A mutant 2B complex**

**Figure S3. Electron density map of ID1 contact**

**Figure S4. Sequence alignment of type I and type II keratins in the 2B region of the rod domain.**

**Table S1. List of ID contacts found in K5/K14-C367A 2B complex**

**Table S2. Summary of the results from PISA and the mutated residues for interaction deficiencies**

**Table S3. Sequences of the oligonucleotide primers used for mutagenesis to produce ID-deficient mutants.**

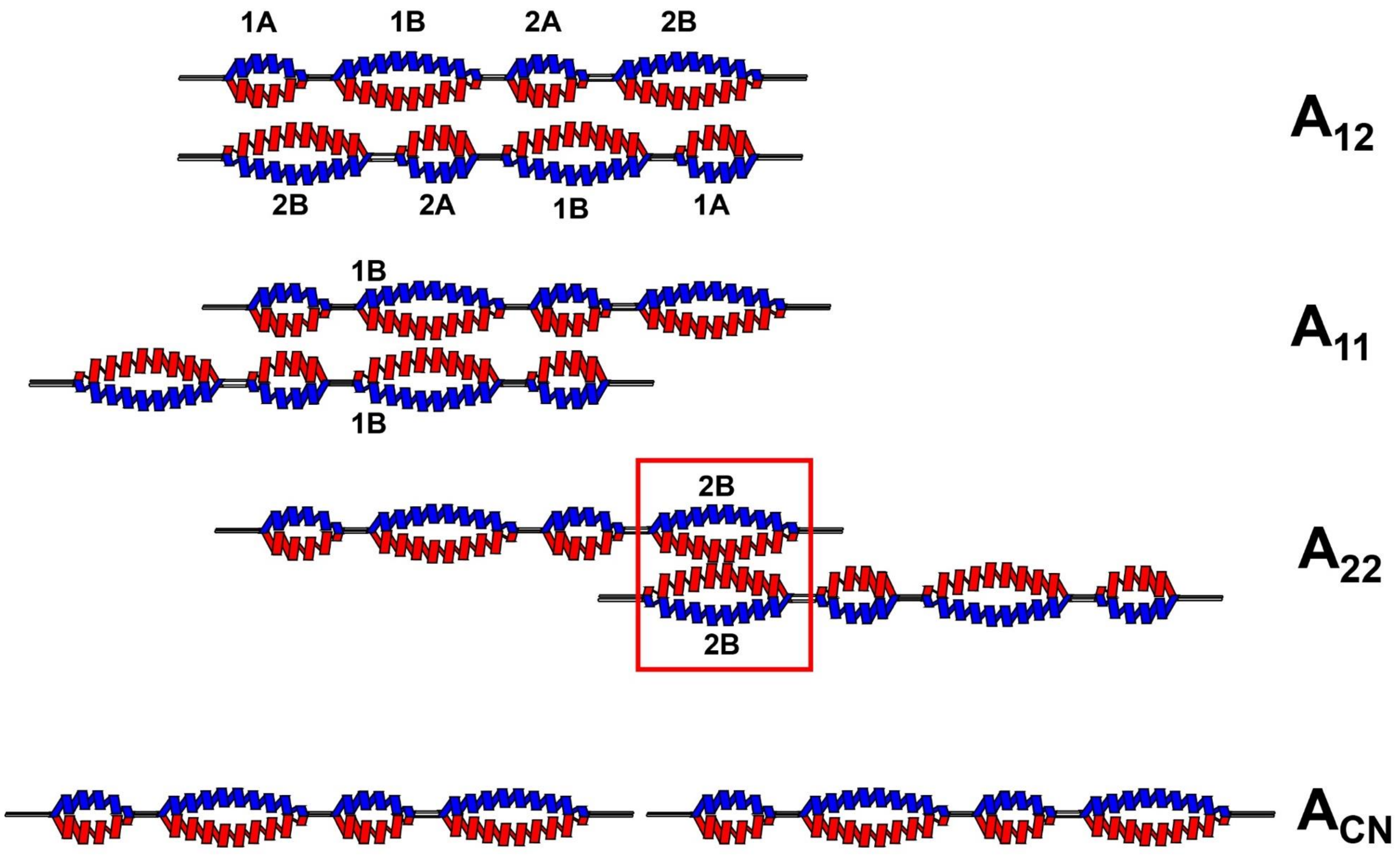


**<u>Figure S1</u>. Four modes of interaction between dimers of IF proteins.** These alignments are named $A_{11}$, $A_{12}$, $A_{22}$, and $A_{CN}$ reflecting the name of the interacting domains involved (Herrmann and Aebi, Curr Opin Struct Biol. 8:177-185. 1998). The interaction modes were suggested from nearest neighbor analysis using chemical cross-linking coupled to mass analysis, complemented by site-directed mutagenesis (Strelkov et al., BioEssays 25:243-251, 2003). The red chains and the blue chains represent type I keratins and type II keratins, respectively.

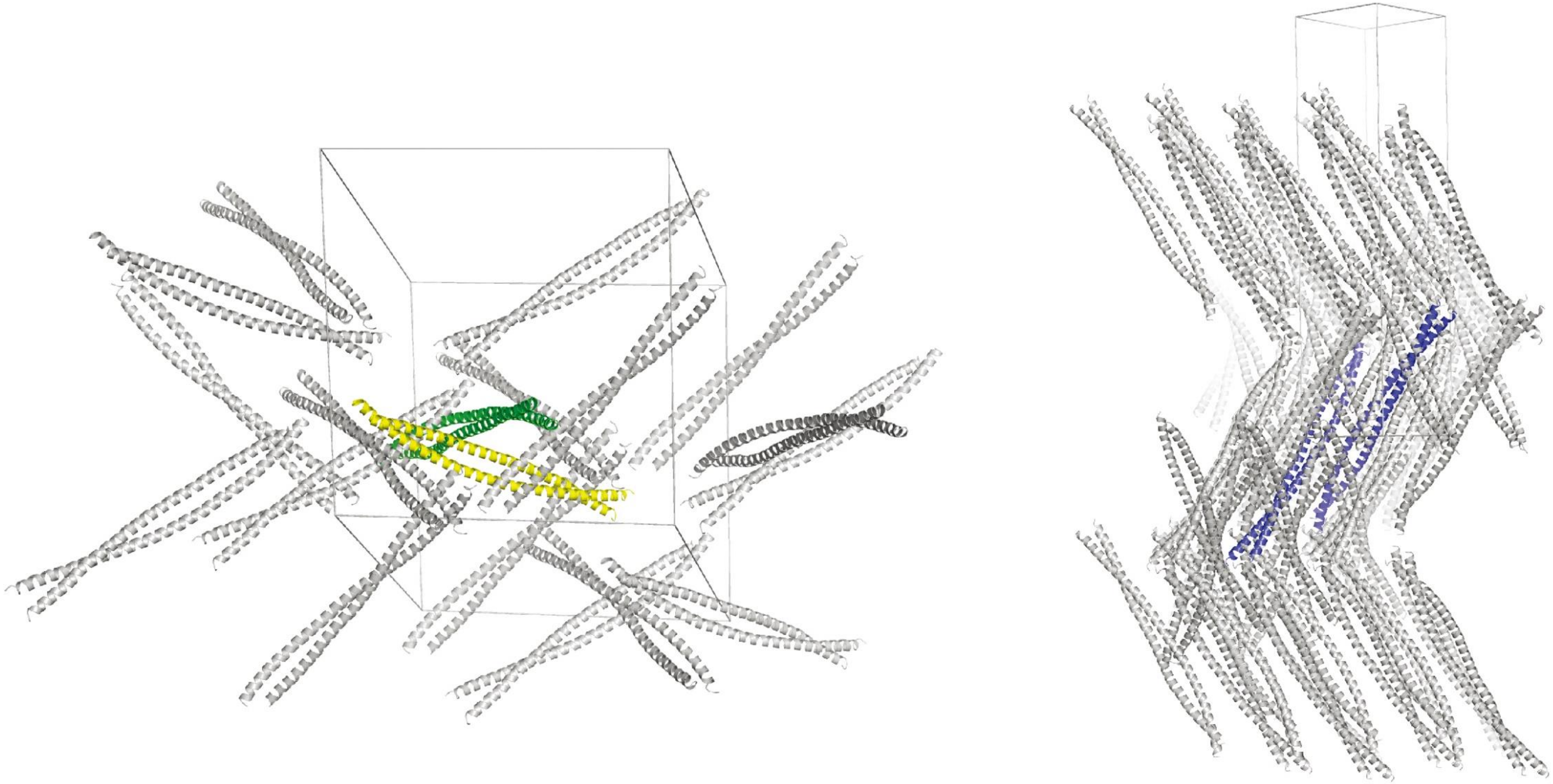

**<u>Figure S2.</u> Crystal contacts of K5/K14 wildtype 2B complex and of K5/K14-C367A mutant 2B complex.** (Left) The crystal contact of wild-type K5/14 2B (PDB ID : 3TNU). the single K5/14 2B heterodimeric coiled-coil helix in asymmetric unit shown in green color and symmetry related molecule making disulfide bond shown in yellow color. (Right) The crystal contact of C367A mutant of K5/14 2B complex (PDB ID: 6JFV). Two heterodimeric molecules in asymmetric unit are shown in blue color. The unit cell is also shown in line.

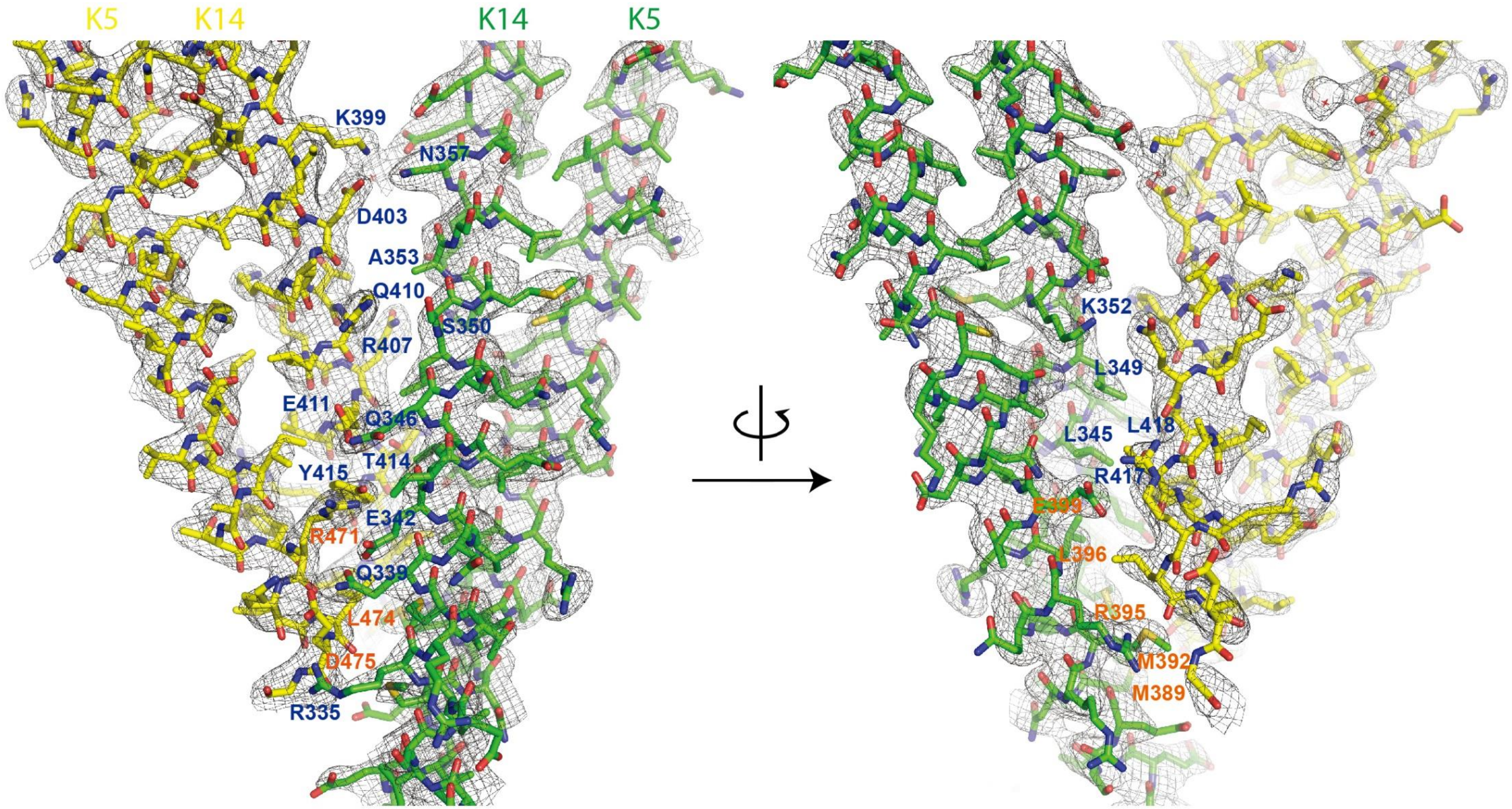


**<u>Figure S3.</u> Electron density map of ID1 contact.** One heterodimeric complex of K14-K5 is shown in green and interacting partner shown in yellow. The residues involved in dimer-dimer interaction shown with residue label. K14 resides are shown in blue and K5 residues are shown in orange.

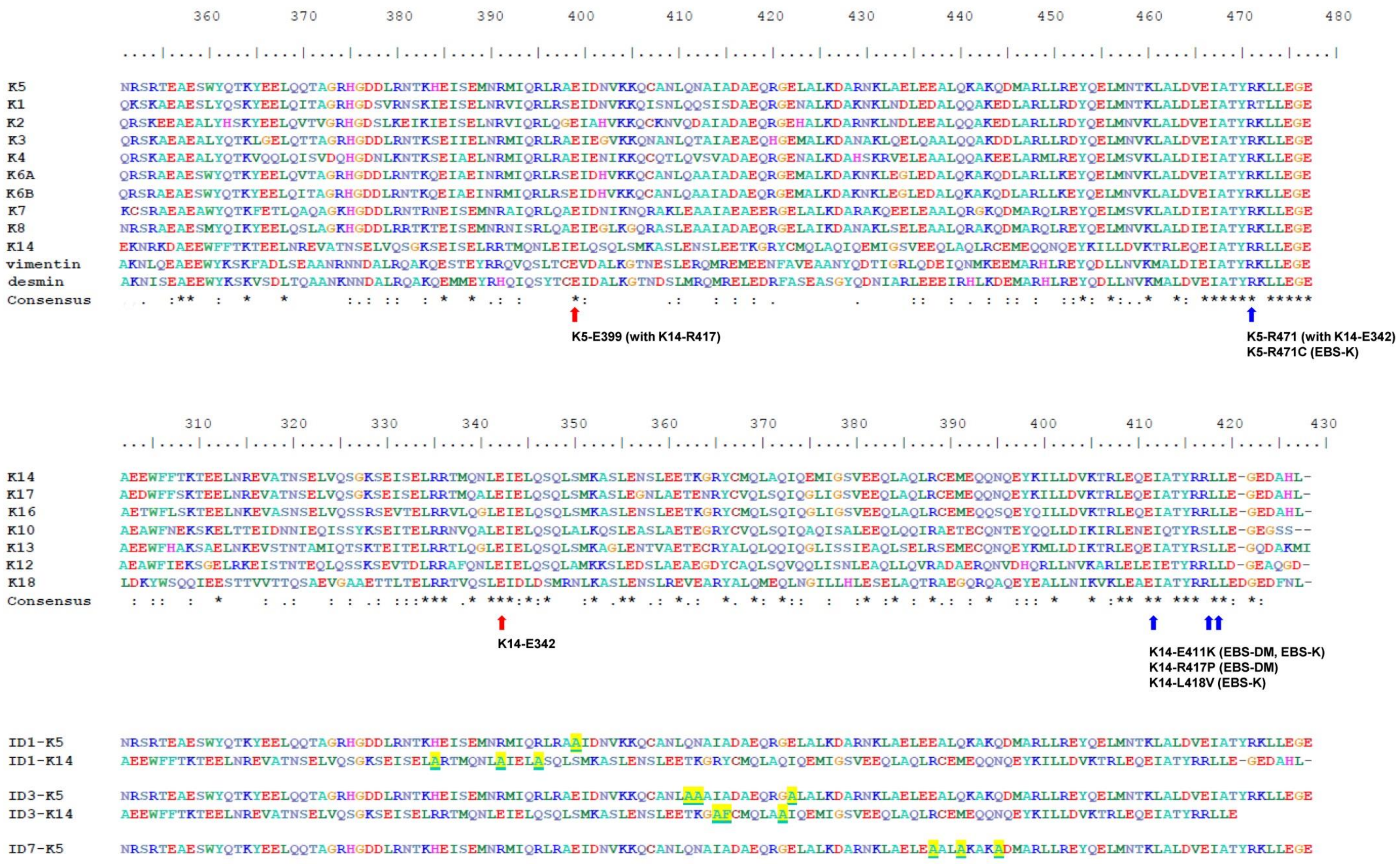


**<u>Figure S4</u>. Sequence alignment of type I and type II keratins in the 2B region of the rod domain.** Note that key amino acids found in both the ID1 and ID3 contacts (Fig. 1) are highly conserved, and some of them are mutated in individuals with EBS. Highlighted and underlined residues are amino acids that were mutated to disrupt dimer interfaces. Red arrows identify key residues not implicated in EBS. Blue arrows identify residues mutated in EBS.

| Contact ID | Interface area ($Å^2$) | Coiled-coil Alignment | Symmetry Operation |
| --- | --- | --- | --- |
| ID1 | 805.7 | Anti-parallel | x, y-1, z |
| ID3 | 760.9 | parallel | -y+1, x, z+1/4 |
| ID5 | 362.3 | T-shape | -y+1, x, z+1/4 |
| ID7 | 215.3 | Anti-parallel | x, y, z |
| ID8 | 190.8 | N/A | -x, -y+1, z-1/2 |
| ID9 | 40.3 | N/A | -y+1, x+2, z+1/4 |
| ID10 | 16.3 | N/A | -x, -y+2, z-1/2 |
| ID11 | 6.5 | N/A | -y+1, x+1, z+1/4 |

**Table S1. List of ID contacts found in K5/K14-C367A 2B complex**

| | ID1 | | ID3 | | ID7 | |
|---|---|---|---|---|---|---|
| Average values | interface area (Å$^2$) | $\Delta^iG$ (kcal/mol) | interface area (Å$^2$) | $\Delta^iG$ (kcal/mol) | interface area (Å$^2$) | $\Delta^iG$ (kcal/mol) |
| | 805.7 | -4.5 | 362.3 | -2.1 | 215.3 | 1.7 |

| dimer 1 | interacting residue (inter-dimer) | Comment | dimer 1 | interacting residue (inter-dimer) | comment | dimer 1 | interacting residue (inter-dimer) | comment |
|---|---|---|---|---|---|---|---|---|
| K14-I331 | - | | K14-M351 | - | | K5-K426 | - | |
| K14-R335 | K5-E475 | mutated to Alanine for deficiency | K14-S354 | K5-Q454 | | K5-N430 | - | |
| K14-M338 | K5-L474 | | K14-L355 | - | | K5-A433 | - | |
| K14-Q339 | K5-E475 | | K14-S358 | K5-Q454 | | K5-E434 | - | |
| K14-E342 | K5-R471<br>K14-Y415 | mutated to Alanine for deficiency | K14-E361 | K5-L461 | | K5-E437 | K5-R451 | mutated to Alanine for deficiency |
| K14-I343 | K5-R471 | | K14-T362 | - | | K5-Q440 | K5-Q444 | mutated to Alanine for deficiency |
| K14-L345 | K14-T414 | | K14-R365 | K5-D464 | mutated to Alanine for deficiency | K5-Q444 | K5-Q440 | mutated to Alanine for deficiency |
| K14-Q346 | K14-E411 | mutated to Alanine for deficiency | K14-Y366 | K5-K460 | mutated to Phenylalanine for deficiency | K5-R448 | - | |
| K14-L349 | K14-Q410<br>K14-E411 | | K14-Q369 | - | | K5-N458 | - | |
| K14-S350 | - | | K14-Q372 | K5-R471 | mutated to Alanine for deficiency | dimer 2 | interaction | note |
| K14-K352 | K14-Q410 | | K5-K404 | - | K404E (WC*) | K5-K426 | - | |
| K14-A353 | K14-R407 | | K5-C407 | - | | K5-N430 | - | |
| K14-E356 | K14-Q410 | | K5-A408 | - | | K5-A433 | - | |
| K14-N357 | K14-D403,<br>K14-K399 | | K5-Q411 | K14-Q394<br>K5-L450 | mutated to Alanine for deficiency | K5-E434 | - | |
| K14-E360 | K14-K399 | | K5-N412 | K14-E397 | mutated to Alanine for deficiency | K5-E437 | K5-R451 | |
| K14-E361 | K14-K399 | | K5-I414 | K5-Y453<br>K5-M457 | | K5-Q440 | K5-Q444 | mutated to Alanine for deficiency |
| K5-E385 | - | | K5-A415 | K14-E397 | | K5-K441 | - | |
| K5-E388 | - | | K5-D416 | - | | K5-Q444 | K5-Q440 | mutated to Alanine for deficiency |
| K5-M389 | K5-E475 | | K5-E418 | K5-K460<br>K14-L401 | | K5-D445 | - | |
| K5-M392 | K5-L474<br>K14-L419 | | K5-Q419 | - | | K5-R448 | - | |
| K5-R395 | K14-E420 | | K5-E422 | K14-R407 | mutated to Alanine for deficiency | K5-R451 | K5-E437 | mutated to Alanine for deficiency |
| K5-L396 | - | | K5-R429 | - | | K5-N458 | - | |
| K5-E399 | K14-R417 | mutated to Alanine for deficiency | dimer 2 | interaction | note | | | |
| K5-V403 | - | | K14-Q394 | K5-Q411 | | | | |
| dimer 2 | interaction | note | K14-E397 | K5-N412<br>K5-A415 | | | | |
| K14-K399 | K14-E360<br>K14-E361 | | K14-I400 | - | | | | |
| K14-D403 | K14-N357 | | K14-L401 | K5-E418 | L401P (WC*) | | | |
| K14-T406 | - | | K14-V404 | - | | | | |
| K14-R407 | K14-A353 | | K14-R407 | K5-E422 | | | | |
| K14-Q410 | K14-E356 | | K14-E411 | - | E411K (DM*, K*) | | | |
| K14-E411 | K14-Q346<br>K14-L349 | E411K (DM*, K*) | K5-M446 | - | | | | |
| K14-T414 | K14-L345 | | K5-A447 | - | | | | |
| K14-Y415 | K14-E342 | Y415H (K*, DM*)<br>Y145C (WC*) | K5-L450 | K5-Q411 | | | | |
| K14-R417 | K5-E399 | R417P (DM*) | K5-R451 | - | | | | |
| K14-L418 | - | L418V (K*) | K5-Y453 | K5-I414 | | | | |
| K14-L419 | K5-M392 | L419Q (DM*) | K5-Q454 | K14-S354<br>K14-S358 | | | | |
| K14-G421 | - | | K5-M457 | K5-I414 | | | | |
| K5-I467 | - | I467L (WC*)<br>I467M (K*)<br>I467T (DM*) | K5-N458 | - | | | | |
| K5-R471 | K14-E342<br>K14-I343 | R471C (K*) | K5-K460 | K5-E418<br>K14-Y366 | | | | |
| K5-L474 | K14-M338 | | K5-L461 | K14-E361 | | | | |
| K5-E475 | K14-R335<br>K14-Q339 | E475K (DM*)<br>E475G (DM*) | K5-D464 | K14-R365 | | | | |
| K5-G476 | - | G476D (WC, EPPK*) | K5-V465 | - | | | | |
| | | | K5-I467 | - | I467L (WC*)<br>I467M (K*)<br>I467T (DM*) | | | |
| | | | K5-A468 | - | | | | |
| | | | K5-R471 | K14-Q372 | R471C (K*) | | | |
| | | | K5-E475 | - | E475K (DM*)<br>E475G (DM*) | | | |

**Table S2**. Summary of the results from PISA and the mutated residues for interaction deficiencies

*Note: Relevance to genetic skin disease is highlighted. Abbreviations are as follow: **DM**, Epidermolysis bullosa simplex, Dowling-Meara variant; **EPPK**, Epidermolytic palmoplantar keratoderma; **K**, Epidermolysis bullosa simplex, Koebner (generalized) variant; **WC**, Epidermolysis bullosa simplex, Weber-Cockayne (localized) variant.

| Mutant | Primer Name | Sequence (5'->3') |
|---|---|---|
| ID1 | ID1_K5E_PvuI_F | CCAGAGGCTGAGAGCCGCGATCGACAATGTCAAGAAACAGTGC |
| | ID1_K5E_PvuI_R | GCACTGTTTCTTGACATTGTCGATCGCGGCTCTCAGCCTCTGGATCA TCCGG |
| | ID1_K14REQ_F | CGAGATCTCGGAGCTCGCGCGCACCATGCAGAACCTGGCGATTGAGCTGGCGTCCC AGCTCAGCATGAAAGC |
| | ID1_K14_K14REQ_R | CCAGGTTCTGCATGGTGCGCGCGAGCTCCGAGATCTCGCTCTTGCCGC TCTGC |
| ID3 | ID3_K5QNE_NotI_F | GCCGAGATTGACAATGTCAAGAAACAGTGCGCCAATCTGGCGGCCGCCATTGCGG ATGCCGAGCAGCGTGGGGCGCTGGCCCTCAAGGATGCC |
| | ID3_K5QNE_NotI_R | GCGGCCGCCAGATTGGCGCACTGTTTCTTGACATTGTCAATCTCGGCTCTCAGCC TCTGGATCATCCGG |
| | ID3_K14RYQ_F | GGAGGAGACCAAAGGTGCCTTCTGCATGCAGCTGGCCGCGATCCAGGAGATGATTGG |
| | ID3_K14RYQ_R | GGCCAGCTGCATGCAGAAGGCACCTTTGGTCTCCTCCAGGCTGTTCTCCAGGG |
| ID7 | ID7_K5AAA_F | GGATGCCAGGAACAAGCTGGCCGAGCTGGAGGCAGCGCTGGCGAAGGCCAAGG CGGACATGGCCCGGCTGCTGCGTGAGTACCAGG |
| | ID7_K5AAA_R | CCTGGTACTCACGCAGCAGCCGGGCCATGTCCGCCTTGGCCTTCGCCAGCGCTGC CTCCAGCTCGGCCAGCTTGTTCCTGGCATCC |

**Table S3. Sequences of the oligonucleotide primers used for mutagenesis to produce ID-deficient mutants.**
